# The pitfalls of regression to the mean in bivariate timeseries analysis

**DOI:** 10.1101/2023.12.02.569707

**Authors:** Thomas M. M. Versluys

## Abstract

1.Plastic traits, capable of taking multiple forms, often correlate with one another or with features of the environment when measured over time. These patterns of correlated change are sometimes assumed to reflect adaptive plasticity, such as coevolved “integrated phenotypes” within individuals, synchronisation between social or mating partners, or responses to environmental gradients.

2. Such plasticity is ecologically and evolutionarily important, so there is considerable interest in understanding how it varies between individuals and groups. However, “regression to the mean”, the statistical tendency for traits to revert to the average value, may create the illusion of strong bivariate correlations in timeseries data, including substantial but meaningless variation between individuals.

3. We demonstrate this using simulated and real data, revealing how regression to the mean can create bias both within and between samples. We then show, however, that its effects can often be eliminated using autoregressive models.

4. We also offer a detailed discussion of how and why regression to the mean arises, introducing the idea that it is both a statistical and ecological phenomenon.

## Introduction

Correlations between two timeseries – that is, sequences of observations measured across time – are commonly interpreted as reflecting stable relationships, where one variable causes corresponding shifts in the other (Bergmüller & Taborsky, 2010; Dingemanse & Araya-Ajoy, 2015). In mated pairs or social partners, synchronised trait changes are seen as efforts to increase social cohesion (Anderson et al., 2003; Bergmüller & Taborsky, 2010; Garcia Castillo et al., 2021; Gray & Ozer, 2019; Laubu et al., 2016; Sjaarda & Kutalik, 2022). In individuals, traits that change in apparent coordination are viewed as indicative of “integrated phenotypes” originating from coevolution (Nielsen & Papaj, 2022; Sheehy & Laskowski, 2023). And when traits seem to respond to changes in the environment, such as weather (Ferderer et al., 2022) or predation risk (Hossie et al., 2017), it is inferred that these short-term adaptations confer some benefit (Fox et al., 2019).

These inferences may often be correct, but “regression to the mean” may also be at work. This is the tendency for plastic traits or those measured with error to converge to the population’s mean value when observed repeatedly (Barnett et al., 2005; Kelly & Price, 2005; Linden, 2013; Yu & Chen, 2015). Under its influence, two variables deviating from their respective means will, on average, trend independently towards them over time. If, when first measured (i.e., at baseline), two variables are concurrently above their means, they will trend downwards and become positively correlated. If they are concurrently below their means, causing them to trend upwards, the same will be true. And if they overlap their means – that is, one is above and one below – they will trend in opposite directions and become negatively correlated. In a dataset of many bivariate timeseries – say, a sample of mated pairs – this process may produce large but misleading variation in patterns of correlated change due to baseline variation alone. If, by chance or design, a sample’s baseline values are systematically biased in one direction, the entire sample may regress to the mean simultaneously, biasing estimates of correlated change between samples. In studies of phenotypic plasticity (Dingemanse et al., 2004; Dingemanse & Araya-Ajoy, 2015; Sheehy & Laskowski, 2023; Stamps & Biro, 2016), differences of this kind could be misinterpreted as evidence that the mechanisms producing plasticity differ when, in fact, they do not.

There is a temptation to view regression to the mean as a purely statistical process, but this is misleading. It is true that, in pre-existing datasets, the process occurs simply because the next observation is more likely to fall around the average value (Barnett et al., 2005; Kelly & Price, 2005). However, if the forces that shaped historical trait distributions – climate, genes, behavioural norms, and so forth – remain intact, future values will follow the same distribution as those past. In such cases, regression to the mean can be thought of as an ecological process that affects new as well as old data. (Fig. 1). In either case, as we show, regression to the mean will manifest as temporal autocorrelation, where a measurement at a given timestep t correlates with its preceding values (e.g., t-1, t-2) or “lags” (Fig. 3; SI Fig. 1).

**Figure. 1.**
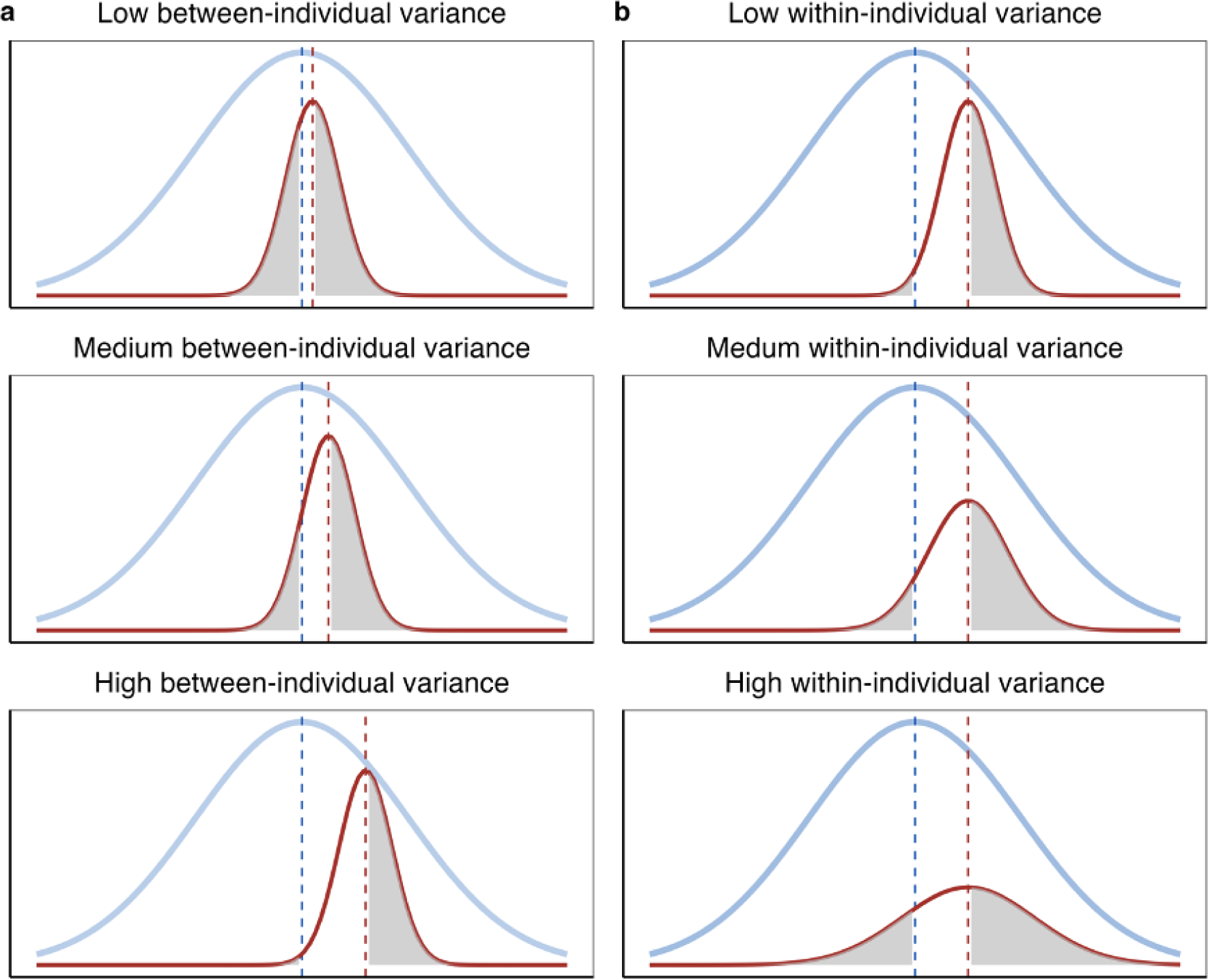
A conceptual view of how data structure may affect population and individual mean regression. Individual distributions (red curves) lie within larger population distributions (blue curves). The grey shaded regions display where regression to the individual mean (red dashed line) will also manifest as regression to the population mean (blue dashed line). In these regions, an apparent trend to the population mean will be in the same direction as a trend to the individual mean.

Whether viewed as a statistical or ecological process, or both, regression to the mean is pervasive in nature. Variables of every kind – from animal abundance (McClure & Rolek, 2023) to body mass (Kelly & Price, 2005) and even stock market returns (Barnett et al., 2005; Kelly & Price, 2005) – come under its influence. Yet it is commonly overlooked and, to our knowledge, has only been studied in univariate timeseries in ecology and evolution (Fournier et al., 2019; Kelly & Price, 2005; McClure & Rolek, 2023). The potential for regression to the mean to bias bivariate studies of correlated change, then, is considerable.

Here, we use simulations and real data from humans, mice, and guppies to demonstrate how regression to the mean affects bivariate studies of correlated change. Drawing on timeseries methods (Class et al., 2017; Dingemanse & Dochtermann, 2013; Houslay & Wilson, 2017; Zuur et al., 2009), we suggest approaches to correct for its effects. We also discuss the circumstances under which regression to the mean is likely to occur and argue that, contrary to the standard description, it makes conceptual and statistical sense to view individuals as regressing to their own individual means, not the population mean.

## Methods

### Data

We used data from humans, mice, and guppies, each measured for a different trait. Our human data included 10 biennial (every two years) body-mass index (BMI) measurements from 1,557 people from the US-based Health and Retirement Survey (HRS). Our mice data was composed of 10 daily records of voluntary wheel rotations from 59 house mice, described in detail elsewhere (Mitchell et al., 2020) and available on Dryad. Our guppy data included daily activity rates for male guppies, recorded as the cumulative distance moved in 20-minute tracking periods over 14 days. The data, originally sourced from three populations at Deakin University, Geelong, Australia, is also described elsewhere (Mitchell et al., 2020) and is available on Dryad. For all analyses, all datasets were grouped by timestep (e.g., “day of measurement” in mice and guppies) and z-standardised (i.e., transformed to have mean zero and variance one). This aided model fitting and removed underlying temporal trends that might obscure the effects of regression to the mean and make comparisons across species difficult. For bivariate analyses, we constructed pairs of timeseries by generating 10,000 random pairings separately for each species. This approach ensured that, on average, bivariate series were uncorrelated in all species, aiding comparison of results.

### Simulation

We also simulated “ground truth” datasets with known properties to test how regression to the mean affects “true” bivariate associations. Our first simulated dataset – hereafter called our “first simulation” – contained 10,000 bivariate timeseries, which, for simplicity, we describe as “pairs” of “individuals”, although they could equally represent traits, animals, or entities of any other kind. Our first simulation was parameterised so that the data had similar properties to our real datasets. Each series had a unique mean drawn from a normal distribution with mean of 0 and variance of 0.3. Subsequent values were generated randomly around this point using an autoregressive (AR1) process with variance of 0.4 and autocorrelation (*rho*) of 0.5, giving each individual a unique distribution of values correlated across time. Individuals within each pair were simulated to be uncorrelated (r = 0) with one another, allowing us to test deviations from the true bivariate association. To investigate the sensitivity of our results to the properties of our data (Fig. 1), we simulated additional datasets, each time modifying one parameter while keeping others constant at the level of our first simulation.

### Statistical analysis

All statistical analysis was performed in the software R (Version 4.2.2). Analyses were performed largely using linear mixed models (R package “nlme”) with covarying slopes and intercepts. We assume, for the sake of simplicity, that the relationships being modelled are linear.

### Do individuals regress to their own means or the population mean?

Before examining the impact of regression to the mean in a bivariate analyses, we wanted to establish how it manifests in a simple, univariate setting. Our ecological perspective leads us to question the traditional idea that individuals regress to the population mean. This is because, in most populations of entities, global causal forces (e.g., climate or shared biology) act alongside local influences that are unique to individuals or groups. Or, alternatively, a causal force might be universal but vary in intensity depending on an individual’s proximity to it. Regardless of how causality is conceptualised, stable phenotypic variation between individuals should emerge, producing many unique trait distributions with different parameters (Barnett et al., 2005). We might expect, ecologically speaking, for each individual to regress to their own distribution’s mean, rather than the population’s mean. We can illustrate this with an example. Consider a population of animals weighing, on average, 30kg. An outlier averaging 50kg in weight and fluctuating between 40kg to 60kg lies on the tail of the distribution. Were this animal measured at 40kg – far above the population mean – we might assume, naively, that it would trend downwards, but this belief would almost certainly be misplaced. The animal, already at the lower bound of its weight, is far more likely to trend upwards towards 50kg than downwards to 30kg.

Had the animal’s individual mean and the population mean been close together (lower between-individual variance), it may indeed have appeared to regress to population mean. Likewise, had the animal’s trait distribution been wider and overlapped the population mean (higher within-individual variance), ranging, say, from 20kg to 60kg, the same would have been true. If we generalise this to all individuals in a sample, it is apparent that the utility of the population mean in predicting individual mean regression depends on the structure of the data (Fig. 1).

However, if a value falls in the white region, this trend to the population mean will be in the opposite direction of the trend to the individual mean. As the white region increases in size due to changes in between-individual (Panel a) and within-individual (Panel b) variance, the expectation that a value should regress to the population mean will become increasingly misleading.

### Humans, mice, and guppies all regress to their own means

We wanted to test whether regression occurs primarily to individual or population means in our ecologically varied datasets. We tested this by first fitting a linear mixed model for each dataset of the form:

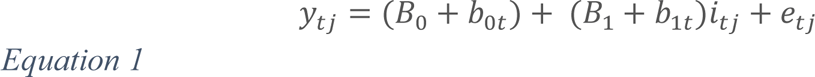

where *y_jt_* is the trait value for *j*th group (in this case, individual) at timestep *t*, *B_0_* is the population intercept, *b*_0j_ and *b*_1j_ are the group-specific intercepts and slopes (i.e., random effects), respectively, *i_tj_* is a continuously coded variable representing time, and *e*0_tj_ is the error term. To derive the trend of each individual over time, we extracted the individual slope estimates from the models, commonly known as “random slopes” or “best-linear unbiased predictors” (BLUPs). We next calculated each individual’s baseline trait (t1) distance from their individual mean (t1 – individual mean) and from the overall population mean (t1 – population mean). These values tell us whether, at baseline, an individual is above or below their own individual and population means, respectively. A positive value indicates a higher-than-mean baseline trait value, while a negative value indicates a lower-than-mean baseline trait value. The magnitude of this value represents the extent of the deviation. Finally, using two separate linear models, we tested which of these distances – individual-mean distance or population-mean distance – better predicted individual time trends. Both models took the form

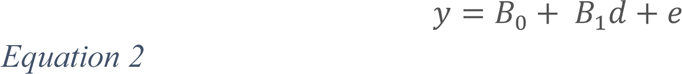

where *y* is the trait value, *B_0_* is the population intercept, *B_1_*is the population slope estimate, *d* is one of the two distance variables, and *e* is the error term.

We found that, in all cases, individual-mean distance explains more than twice the slope variance of population-mean distance and has a better model fit, as measured by ANOVA with a Chi-squared test (Table 1).

**Table. 1.** Comparison of model fits for predicting individual trends based on deviation from population means and individual means. Two goodness-of-fit statistics (r² and residual sum of squares, RSS) are presented for the “population-mean model” and the “individual-mean model”. These models were used to predict individual trends in traits across humans, mice, and guppies for body-mass index (BMI), wheel rotations, and daily movement distance, respectively. Higher r² and lower RSS values indicate better model fit.

|  |  |  | Population-mean model |  | Individual-mean model |  |
| --- | --- | --- | --- | --- | --- | --- |
| *Species* | *Trait* | *Measurement frequency* | $r^2$ | RSS | $r^2$ | RSS |
| Humans | Body-mass index | Biennial (every two years) | 0.08 | 273 | 0.48 | 153.1 |
| Mice | Wheel rotations | Daily | 0.07 | 12.3 | 0.8 | 2.7 |
| Guppies | Movement distance | Daily | 0.17 | 94.6 | 0.42 | 66.5 |

Models using BLUPs should be interpreted cautiously, as uncertainty (e.g., standard errors around slope estimates) is not propagated through to subsequent analyses (Houslay & Wilson, 2017), but these results give a strong impression that regression to the mean is typically an individual-centric process. However, this may not always be true, so each dataset should be examined individually.

Having established that regression to the mean is an individual-centric process in humans, mice, and guppies, we then refit our models (Equations 1 and 2) in simulated datasets to test how data properties affect the relative importance of individual and population means. The relative performance of the population-mean model, as predicted (Fig. 1), declined with higher between-individual variance and improved with higher within-individual variance (Fig. 2).

**Figure 2.**
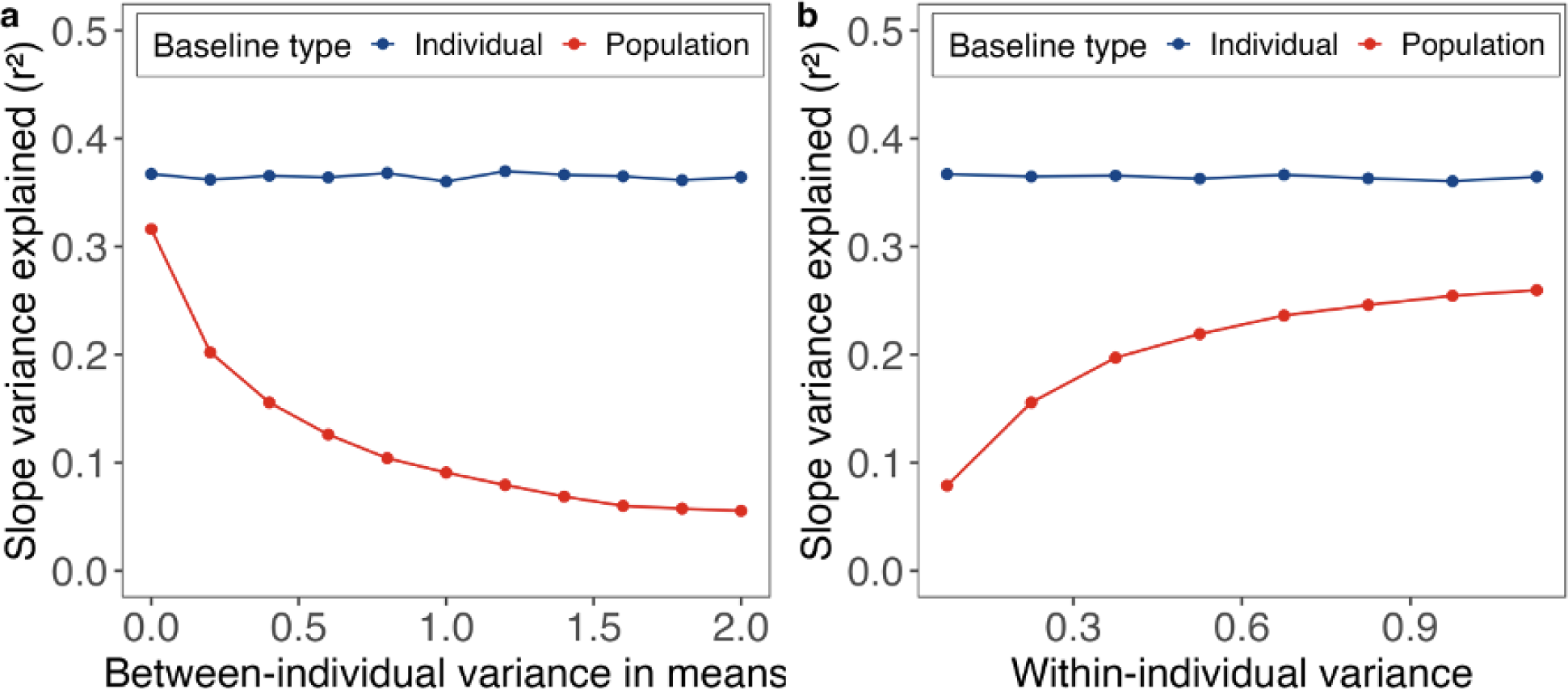
A comparison of population and individual means as predictors of linear time trends. The proportion of the variance in time trends (random slopes) explained by baseline distance from individual means (blue) and baseline distance from the population mean distance (red), as a function of the between-individual variance in means (Panel a) and the within-individual variance (Panel b). Results are calculated as the average from identical models fit in 100 simulated datasets.

### Regression to the mean biases estimates of correlated change

Having established that regression to the mean occurs and, in our datasets, is an individual-centric process, we next investigated its impact on bivariate associations. We did so initially in simulated datasets where the properties of the data were carefully controlled. Once again, we fit linear mixed models with covarying slopes and intercepts, this time regressing one individual’s series on their partner’s series in a bivariate model of the form

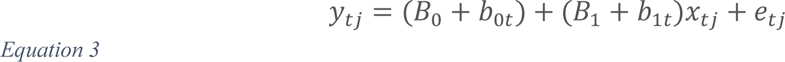

where *y_tj_* and *x_tj_* are the trait values for Individuals 1 and 2 for *j*th pair (group) at observation *i*, *B*_0_ and *b*_0t_ are population intercept and slope, respectively, *b*_1_ and *b*_1t_ are the group-specific intercepts and slopes, respectively, and *e*0_tj_ is the error term. We fit this model in 100 simulated datasets and averaged across the results, repeating this process for low (0.2), medium (0.4), and high (0.6) autocorrelation datasets to explore the impact of autocorrelation.

To visualise the effects of bivariate regression to the mean, we extracted random slopes for each pair and plotted these against the pair’s joint baseline distribution (Fig. 3). Before calculating baselines, we grouped data by individual and z-standardised within each group, so that each individual’s mean was 0 and their standard deviation was 1. This allowed baselines to be interpreted as the distance in standard deviations from individual means, which, as already established, are stronger predictors of time trends that the population mean (Fig. 2). We then took the first of these observations as an individual’s baseline. Within-individual standardised values were only used to calculate baselines and not as the predictors or response variables in any models.

**Figure 3.**
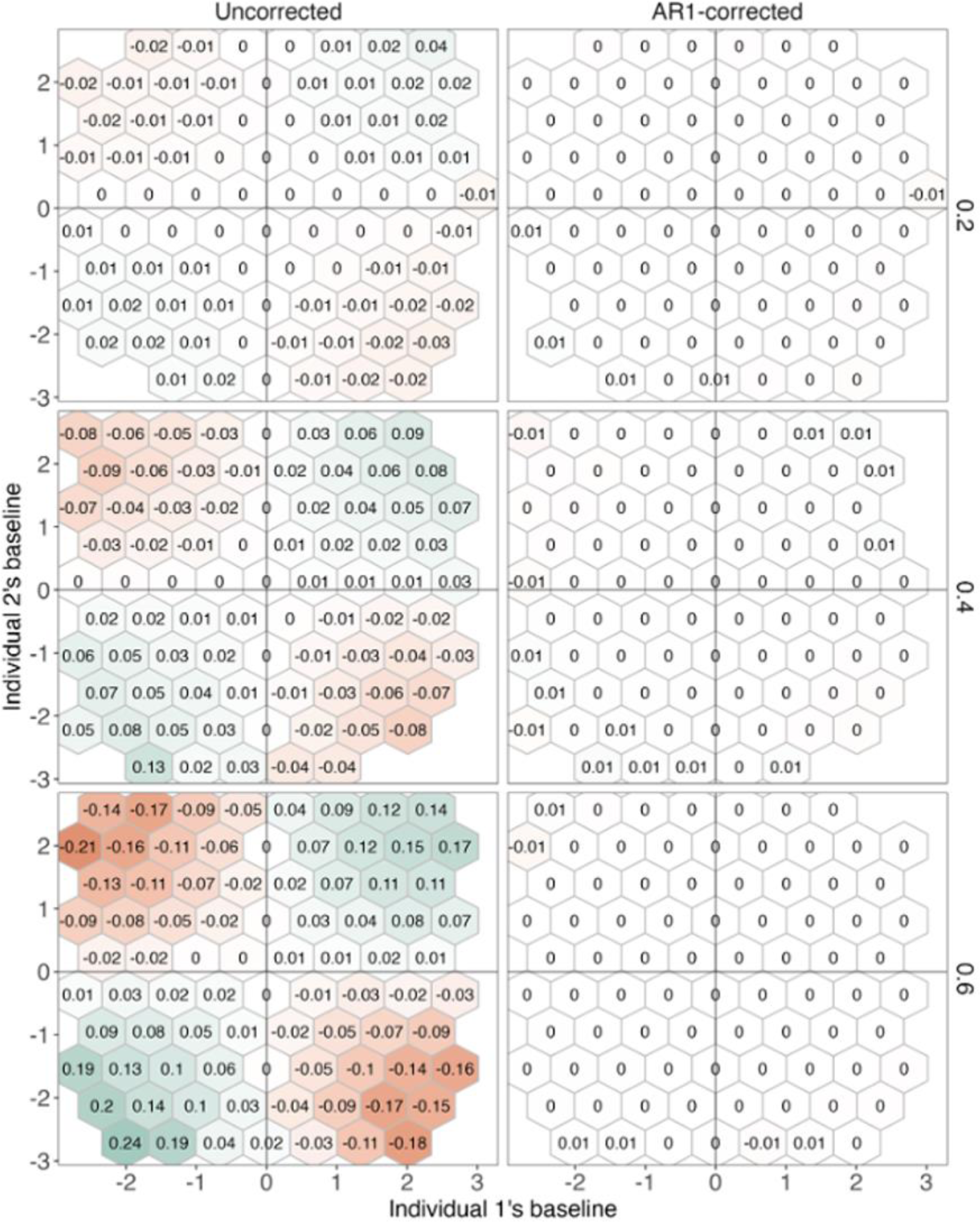
The impact of regression to the mean on bivariate slope estimates at different levels of autocorrelation. Average slopes estimates from 100 simulations are shown for uncorrected (left panels) and AR1-corrected (right panels) models at autocorrelation (rho) of 0.2, 0.4, and 0.6. The x and y axes display the baselines of each individual, scaled within-individual so that baselines can be interpreted as an individual’s distance in standard deviations from their own means. The hexes corresponds to the mean estimated slope for pairs falling within a particular joint baseline area. Colours indicate the magnitude of negative (red) or positive (green) slopes. The central vertical and horizontal lines in each plot represent the “zero” baseline levels.

We found that regression to the mean has a strong impact on a pair’s bivariate association (Fig. 3). If, at baseline, pairs overlapped their respective means, they tended to have negative slopes, whereas those simultaneously above or below their means had positive slopes. Slope estimates decayed towards zero (the “true” simulated value) as one or both baselines moved closer to the mean (Fig. 3a). The strength of these effects, as expected, increased markedly as autocorrelation rose from 0.2 to 0.6, as did random slope variance (SI Fig. 2), which, across the autocorrelation range, rose by 188% from 0.08 to 0.23.

We reasoned that since regression to the mean appears to be an autoregressive process, it could be eliminated with a first-order autoregressive (AR1) model of the form

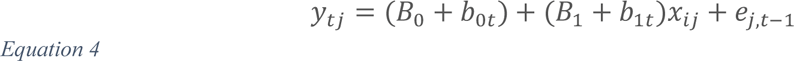

where all else remains the same as in Equation 3 but *e*_j,t−1_ represents the error term being regressed on itself. This model accounts for temporal autocorrelation in the model’s residuals and adjusts outputs accordingly, assuming that the strength of autocorrelation falls with distance between observations (Zuur et al., 2009). Our AR1 model eliminated all baseline dependence (Fig. 3), regardless of the level of autocorrelation, and shrunk slope variance to approximately zero (SI fig. 2). In our simulated data, therefore, an autoregressive model is an effective way to control for regression to the mean.

### What is the impact of sample size and series length?

Since datasets often vary in series length (i.e., number of timesteps) or sample size (i.e., number of pairs), we next investigated whether our results depended on these features of our data. To do so, we adjusted series length from 10 to 50 and sample size from 2,000 to 20,000, holding other parameters at original levels, refit our models, and, as before, repeated this process 100 times. We found that, as expected, baseline dependence is weaker in longer series (Fig. 4; SI Fig. 2), but is still present in series 50 timesteps long, which exceeds the length of many ecological series. We did not find that sample size had any effect on parameters of interests (SI Fig. 2).

**Figure 4.**
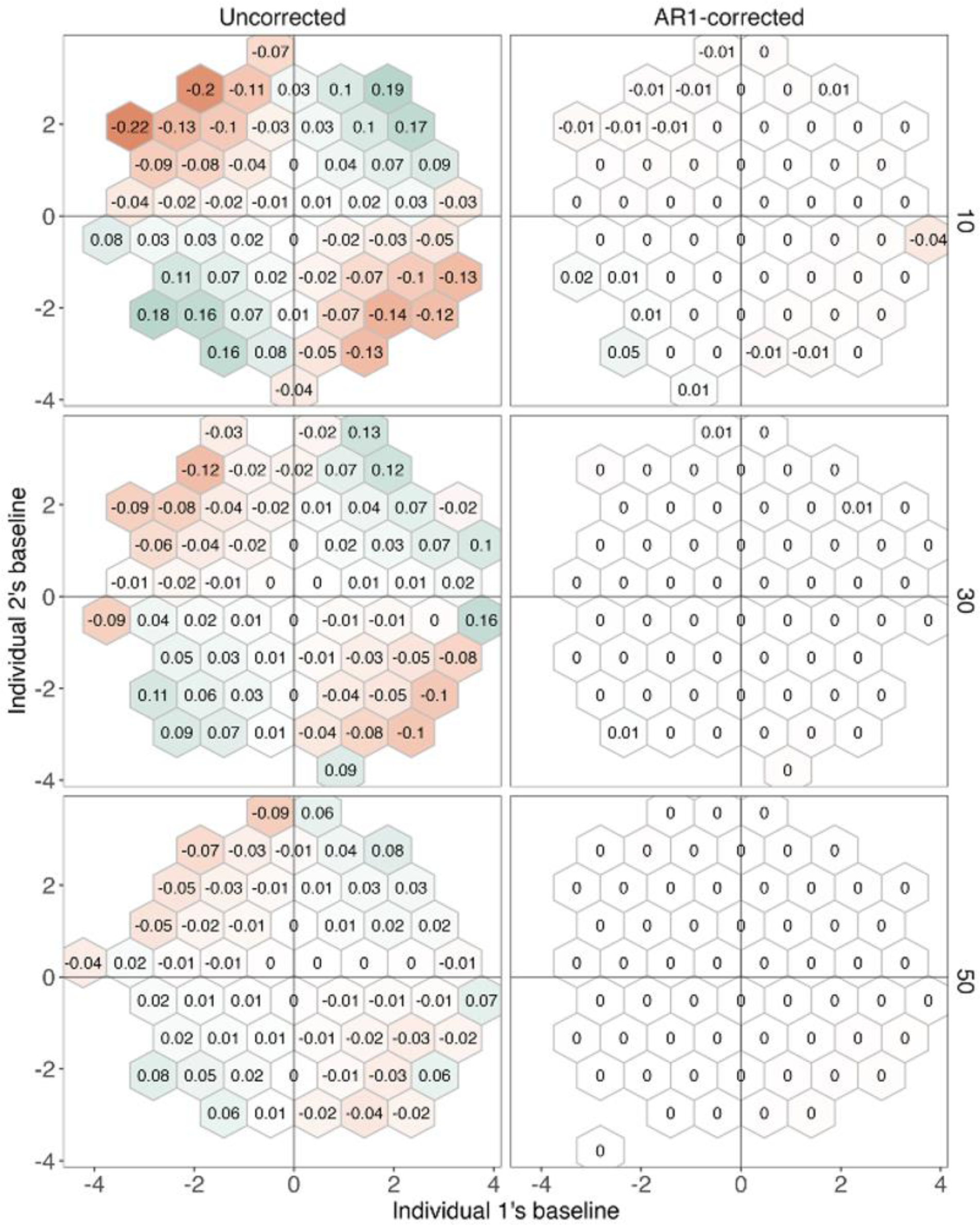
The impact of regression to the mean on bivariate slope estimates at different series lengths. Slopes estimates, averaged across 100 simulations, are shown for uncorrected (left panels) and AR1-corrected (right panels) models at series lengths of 10, 30, and 50. The x and y axes display the baselines of each individual, scaled within-individual so that baselines can be interpreted as an individual’s distance in standard deviations from their own means. The hexes corresponds to the mean estimated slope for pairs falling within a particular joint baseline area. Colours indicate the magnitude of negative (red) or positive (green) slopes. The central vertical and horizontal lines in each plot represent the “zero” baseline levels.

### Bias affects entire samples and is harder to resolve

An individual’s position relative to their mean, then, affects their time trend, which in turn strongly influences bivariate associations in pairs. But what about situations in which, by chance or design, a large proportion of samples, or even all of them, are regressing to the mean simultaneously? To investigate this, we introduced sampling bias into our data, creating two datasets where: i) when a pair’s baselines, defined in the same way as before, are both 0.5 SD above or below the mean; and ii) when a pair’s baselines both overlap the mean, such that one individual is > 0.5 SD above and the other is < −0.5 SD below. We fit our models on both datasets, extracted random slopes, and plotted these against baselines.

We found, as expected, that slope estimates in all individuals were biased in a direction determined by baselines, but, unlike before (Fig. 3), this was also true in AR1 models (Fig. 5a). This may be because these models evaluate autocorrelation from the relationship between consecutive residuals (Zuur et al., 2009) and, when all individuals trend in one direction, there is less dependence between sequential residuals, causing autocorrelation to be underestimated. We can see this by inspecting the autocorrelation parameter (*phi*) in the AR1 models. In our model fit on normal “bias-free” data, *phi* is estimated at the simulated level of 0.5, whereas it is estimated at 0.14 in both models fit on systematically biased data. We are not aware of a solution to this problem.

**Fig. 5:**
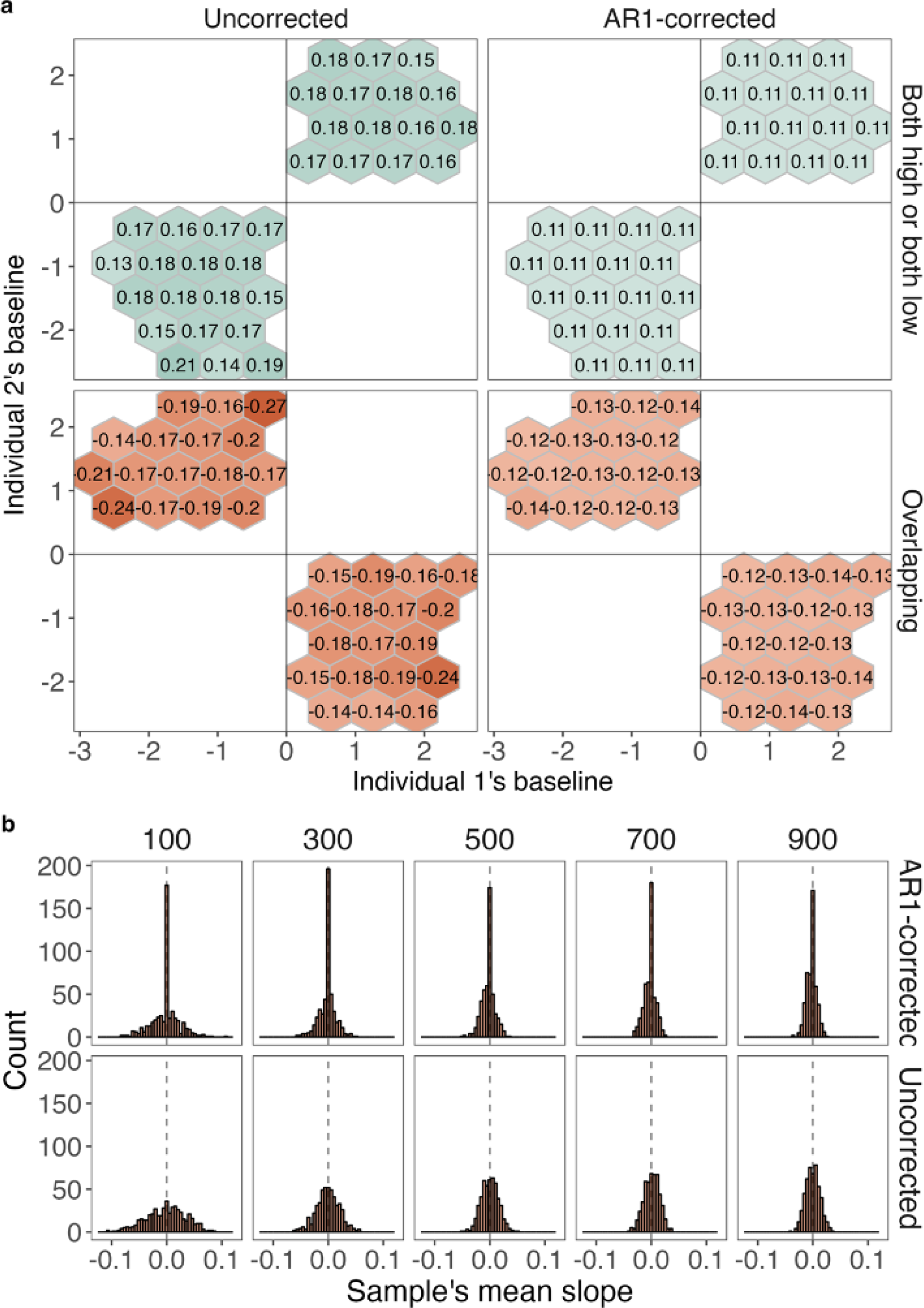
Regression to the mean due to sampling bias. Panel a displays slope estimates for uncorrected (left panels) and AR1-corrected (right panels) models in samples where all pairs are either on the same side (Both high or both low) or different sides of the mean. The x and y axes display the baselines of each individual, scaled within-individual so that baselines can be interpreted as an individual’s distance in standard deviations from their own means. The hexes corresponds to the mean estimated slope for pairs falling within a particular joint baseline area, averaged across 100 simulations. Colours indicate the magnitude of negative (red) or positive (green) slopes. The central vertical and horizontal lines in each plot represent the “zero” baseline levels for each individual. Panel b displays the variance in mean slope estimates (*B)* in 500 samples, ranging in size from 100 to 900 pairs. Each bar represents the count of mean slopes for a particular range, with the dashed vertical line representing a slope estimate of zero. The histograms are grouped by model type (Uncorrected or AR1-corrected) on vertical axis and sample size (100 to 900 pairs) on the horizontally.

Should this be a cause for concern? In many circumstances, probably not. We found that if we took 500 samples of our data, each just 100 pairs in size, fit bivariate models and took the mean slope estimate (*B*) for each sample, the risk of drawing a systematically biased sample at random was extremely low (Fig. 5b). Across 500 samples, the standard deviation in mean slope estimates was just 0.03, with a 96% chance of sample’s mean slope falling within the range −0.06 to 0.06. A sample of just 100 pairs, of course, is small and we found that slope variance declined further if we increased sample size (Fig. 5b), falling to just 0.013 in samples 900 pairs in size.

Slope variance was still lower in AR1 models, at just 0.02 and 0.008 in the 100-pair and 900-pair scenarios, respectively (Fig. 5b). In reality, sampling may not be random, as similar individuals often cluster together in structured populations (Leslie et al., 2015; Rudolf & Rasmussen, 2013), so the results presented here may underestimate the risk of systematic bias.

### Case studies in humans, mice, and guppies

In real datasets of humans, mice, and guppies we found strong parallels with our simulated data. There was strong baseline-dependence in all cases, illustrating that regression to the mean significantly impacts bivariate associations for varied traits in multiple species. As in simulated data, pairs at their respective means typically had negative slopes, while those simultaneously above or below their means had positive slopes. As one or both baselines shifted closer to the mean, slope estimates tended towards zero (Fig. 6). When we used AR1 models, baseline-dependence largely disappeared. Random slope variance was notably higher in uncorrected models for humans, mice, and guppies (sd = 0.27, 0.13, and 0.25, respectively) when compared to AR1 (sd = 0.06, 0.05, and 0.05, respectively). However, mean slope estimates were largely unaffected by regression to the mean, remaining stable in humans, mice, and guppies in uncorrected (*B* = 0, −0.02, and 0.03, respectively) and AR1 (B = 0, −0.02, and 0.01, respectively). However, despite using random pairs, slopes were not exactly zero, implying the presence of residual structure in the data models. As in simulated data, we found that in samples with systematic baseline bias (high/low or overlapping), AR1 models were only partly able to rectify the issue (SI Figs. 3-5)

**Figure 6.**
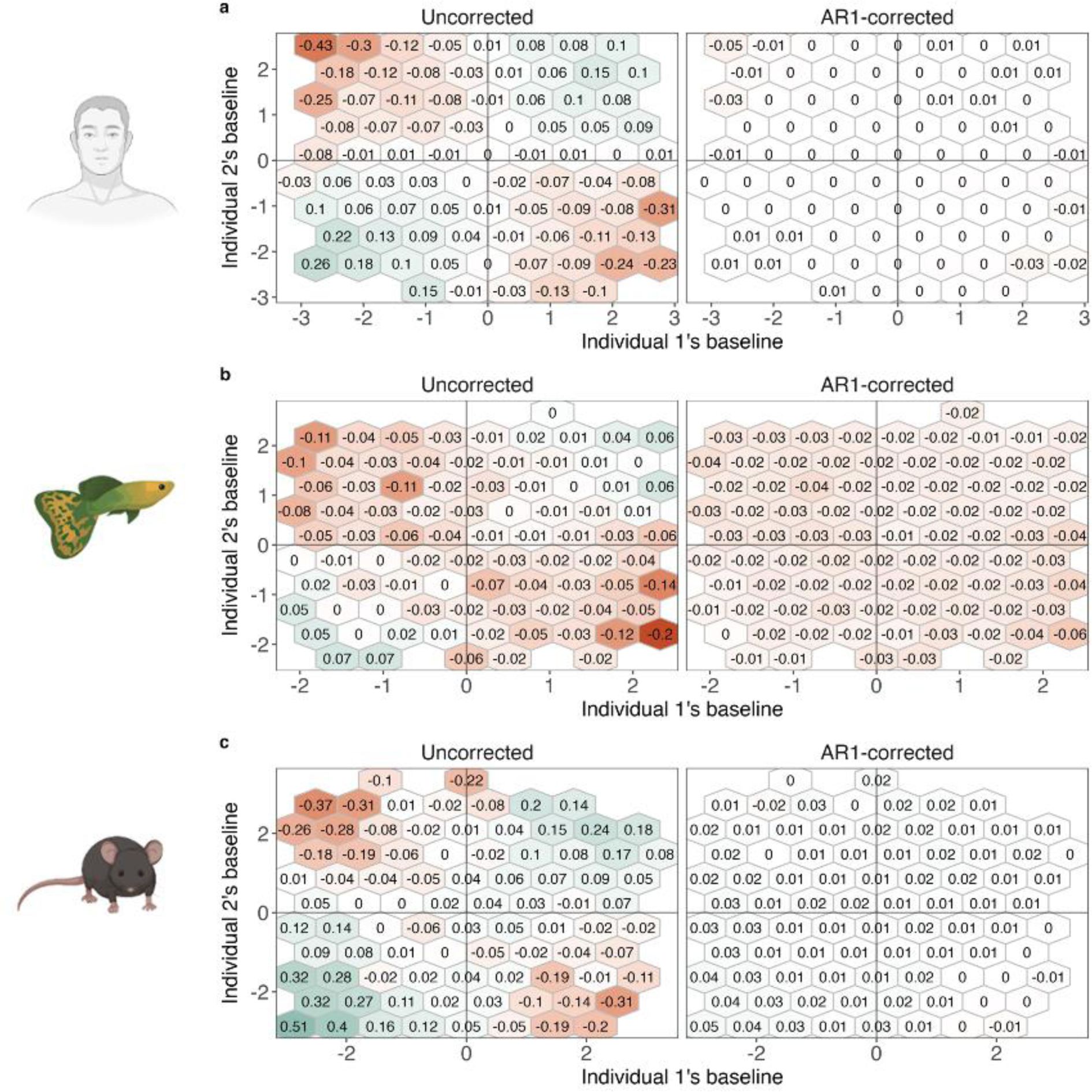
The impact of regression to the mean on bivariate slope estimates in real data. Results are shown for humans (a), mice (b), and guppies (c) in uncorrected (left panels) and AR1-corrected (right panels) models. The x and y axes display the baselines of each individual, scaled within-individual so that baselines can be interpreted as an individual’s distance in standard deviations from their own means. The hexes corresponds to the mean estimated slope for pairs falling within a particular joint baseline area. Colours indicate the magnitude of negative (red) or positive (green) slopes. The central vertical and horizontal lines in each plot represent the “zero” baseline levels.

## Discussion

Regression to the mean causes substantial bias when studying associations in bivariate timeseries. This is true in humans, mice, and guppies for diverse traits measured at very different time scales, indicating that the problem affects ecological data of many kinds. Since these bivariate timeseries associations are used to infer trait plasticity in a variety of study systems (McGuigan et al., 2011; Sheehy & Laskowski, 2023; Stamps & Biro, 2016), the biases we observe here could either obscure genuinely meaningful findings or lead to spurious inferences. If biased slope estimates, which often represent fleeting, meaningless variation, are used to categorise individuals or pairs into groups (Houslay & Wilson, 2017), which his relatively common practice (Laubu et al., 2016), downstream analyses might also be affected.

This bias likely extends beyond the issues discussed here. For example, if individuals positioned on opposite sides of their respective means become in absolute terms, this may create the impression of deliberate trait convergence – that is, the movement of traits towards each other – in mated pairs or other social partners. This may be interpreted as an adaptive strategy to increase similarity, as it has been species ranging from mosquitos (Aldersley et al., 2016; Garcia Castillo et al., 2021) to humans (Anderson et al., 2003; Sjaarda & Kutalik, 2022).

We cannot rely on sample size or series length to mitigate bias arising from regression to the mean. When bias was judged in terms of baseline dependence, sample size had no effect (SI Fig. 2), while series length was not able to eliminate the effects of regression to the mean even by 50 timesteps (Fig. 4). A solution in both simulated and real data appears to be AR1 models, which eliminated the effects of regression to the mean almost entirely (Fig. 1a-b). This supports our claim that the phenomenon is a specific case of temporal autocorrelation and provides a simple solution to address for it in bivariate analyses. More complex autoregressive models may sometimes be required (Harrison, 2021; Zuur et al., 2009), but this was not true in our data.

We were not able to fully resolve the problem of systematic bias that arises when a large proportion of a sample regresses to the mean simultaneously (Fig. 5a). Our simulation of a random sampling scenario indicates that such bias is unlikely to arise by chance (Fig. 5b), but, in practice, the natural structure of populations means that sampling will rarely be random. It is easy to envision how structure could introduce bias into a sampling process. Spatial bias, for example, could be introduced if animals sampled from a resource-rich food patch have unsustainably high baseline body weights, leading to a sample-wide decline thereafter. Temporal bias could occur if samples are collected on a particularly hot day, producing unusual baseline physiological parameters that normalise over time (Naya et al., 2018). Such risks may be best mitigated by awareness at the sampling stage of research.

An underlying message of our work is that regression to the mean is deeply intertwined with ecology and evolution. Even as a purely statistical phenomenon, it represents the influence of previous biological processes that acted on historical populations and now manifest in our datasets. These forces may linger on, shaping contemporary or future populations and the datasets that represent them. A clearer understanding of these forces will help us understand how, why, and where our data is trending.

## Supplementary materials

**SI Fig. 1.**
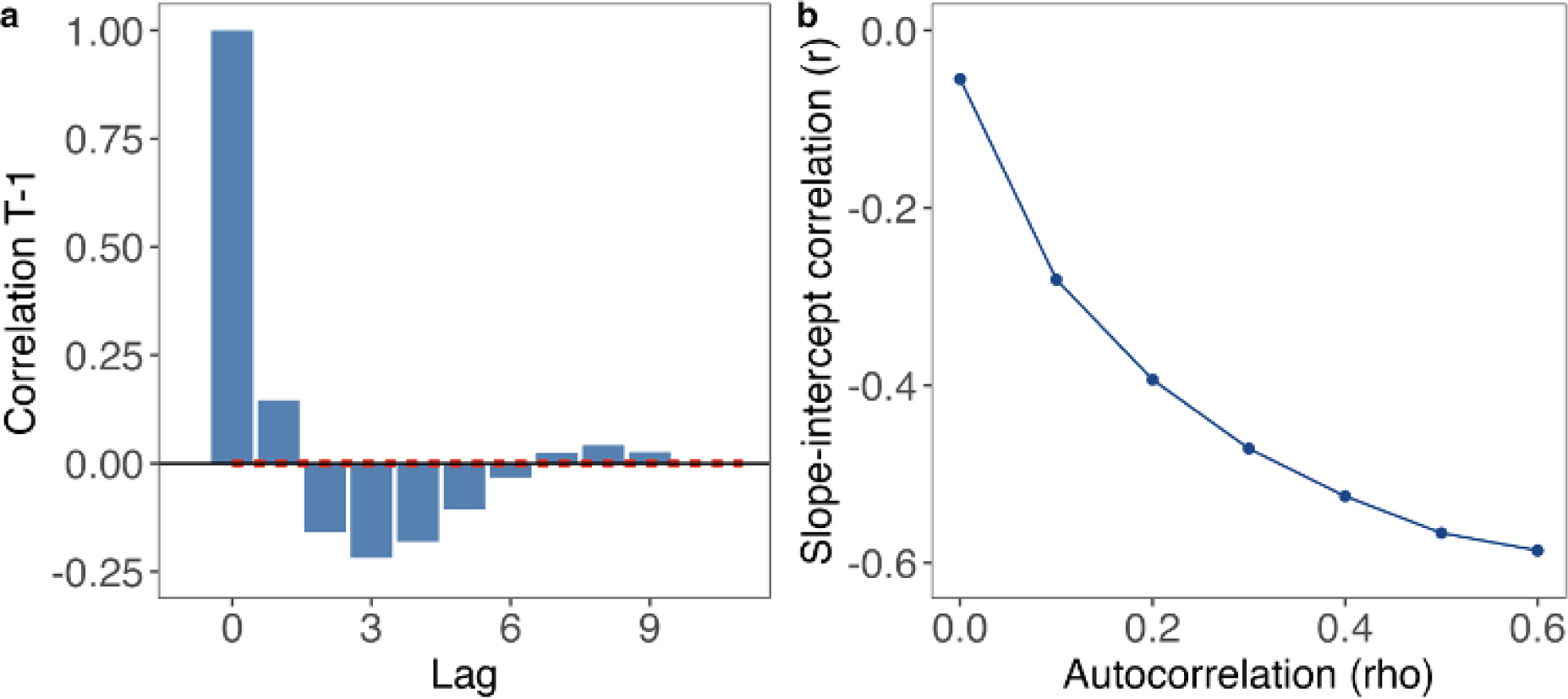
Regression to the mean in univariate timeseries. Panel a shows an autocorrelation plot displaying the correlation between timesteps (Lag) at increasing distances in our first simulated dataset of 10,000 pairs. Lag 0, for example, displays the correlation of a timestep with itself; lag 1 the correlation with values one timestep away, and so forth. Panel b displays the slope-intercept correlation at different levels of temporal autocorrelation, estimated from the random effects structure of linear mixed models (100 iterations for each rho) testing trait scores over time. If regression to the mean is occurring, individuals with high intercepts should trend down (negative slope) and those with low intercepts should trend up (positive slope), leading to a negative slope-intercept correlation. This, indeed, is what we found (r = −0.56), but, consistent with the idea that regression to the mean is autocorrelation, we found that the slope-intercept correlation depended almost entirely on the level of temporal in our simulation, approaching zero as autocorrelation disappears.

**Fig. S2.**
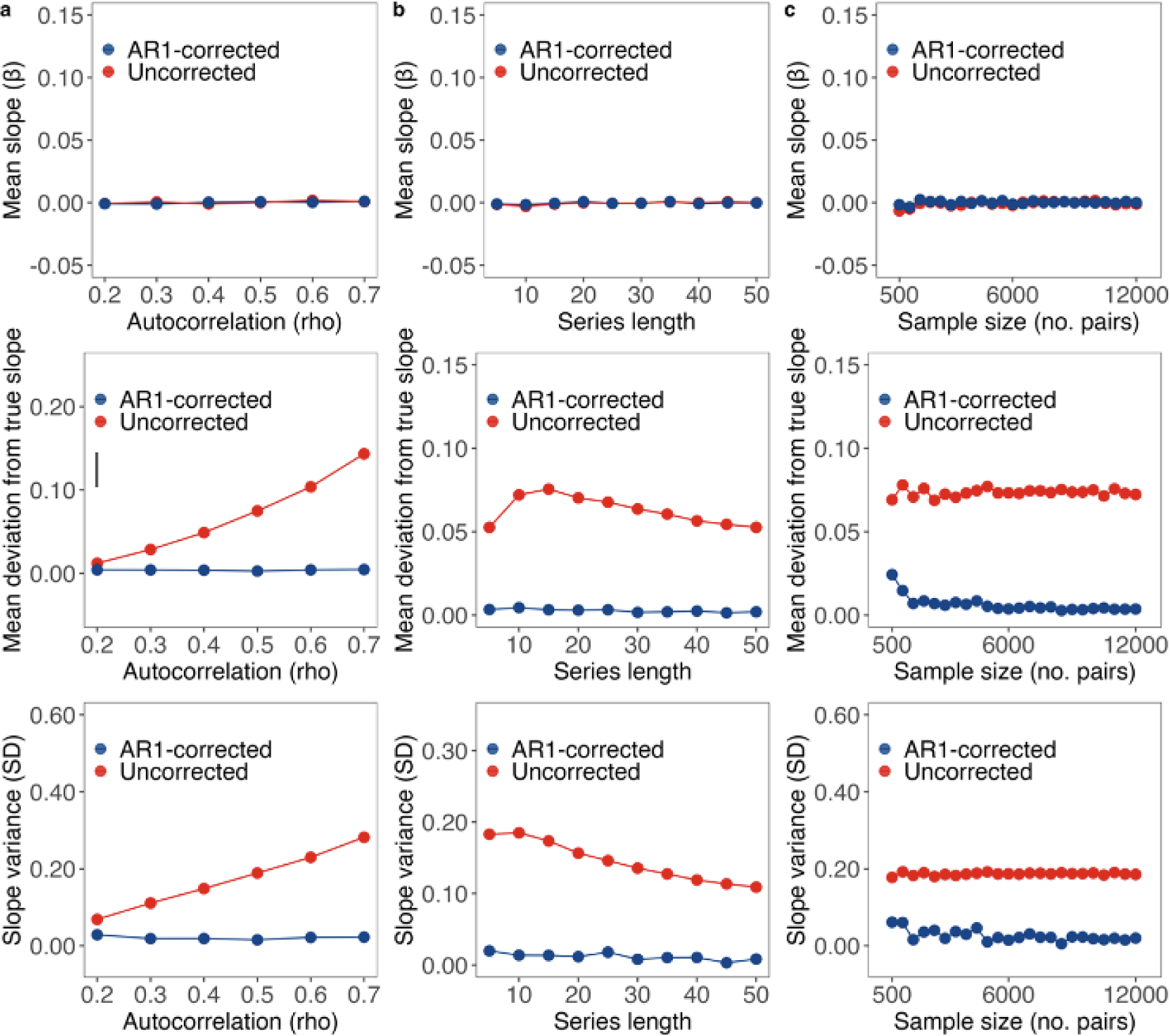
We generated a large number of new datasets, each time modifying a specific parameter on a continuum from high to low, while keeping all other parameters constant at the level of our initial simulation. For each newly simulated dataset, we refit our uncorrected and AR1-adjusted bivariate models, and examined output metrics often used in studies of correlated change. We repeated this process 100 times, each time calculating the average of three specific outputs: i) the average slope estimate or the fixed effect (β); ii) the random slope variance (u), quantified in standard deviations and derived from the model’s random effects; and iii) the average deviation from the mean slope, which can be viewed as a measure of bias, and was determined by extracting each individual’s slope estimate, computing the deviation from the “true slope” of zero, and then averaging these differences.

**SI Fig. 3.**
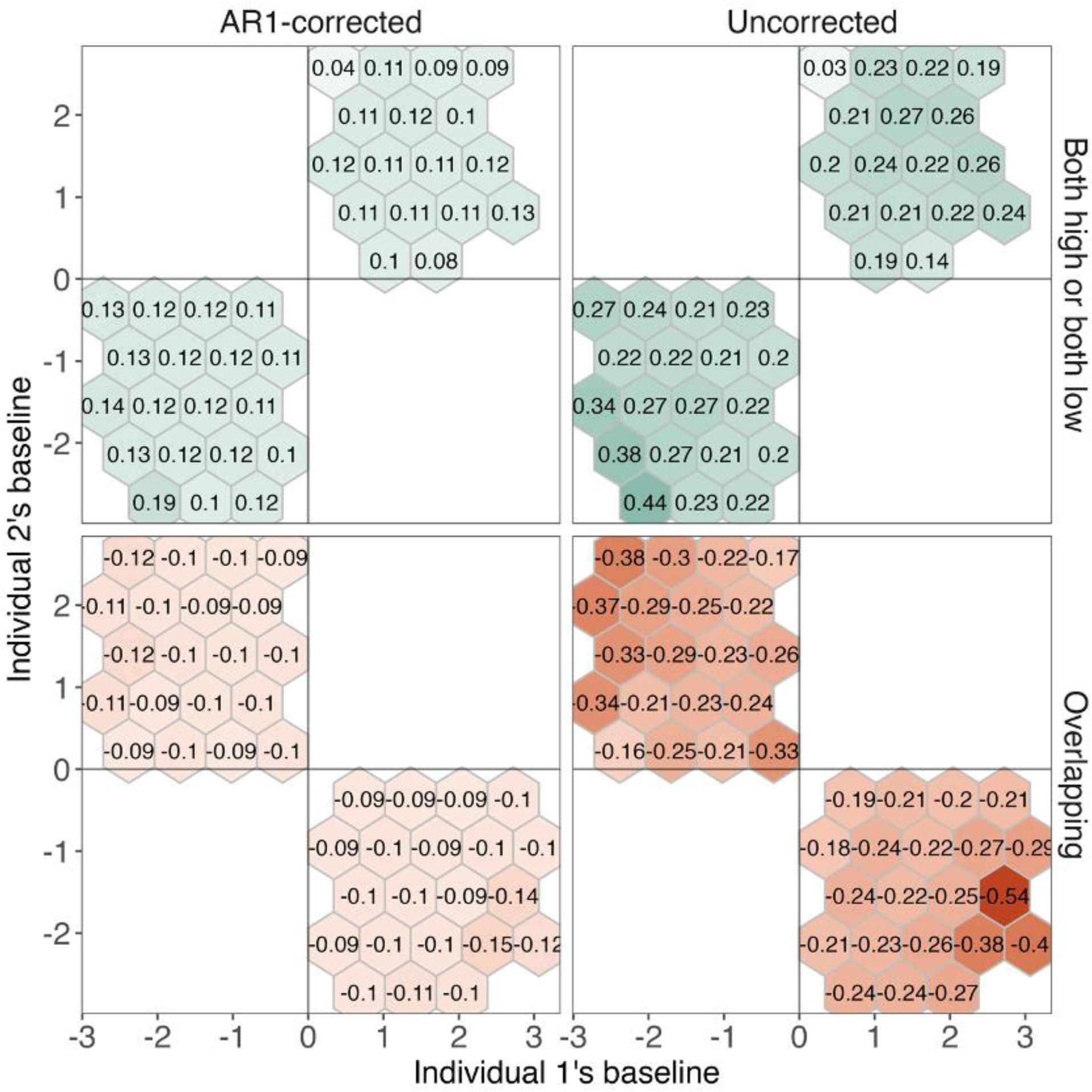
Regression to the mean due to sampling bias in humans. Panel a displays slope estimates for uncorrected (left panels) and AR1-corrected (right panels) models in samples where all pairs are either on the same side (Both high or both low) or different sides of the mean. The x and y axes display the baselines of each individual, scaled within-individual so that baselines can be interpreted as an individual’s distance in standard deviations from their own means. The hexes corresponds to the mean estimated slope for pairs falling within a particular joint baseline area. Colours indicate the magnitude of negative (red) or positive (green) slopes. The central vertical and horizontal lines in each plot represent the “zero” baseline levels for each individual.

**SI Fig. 4.**
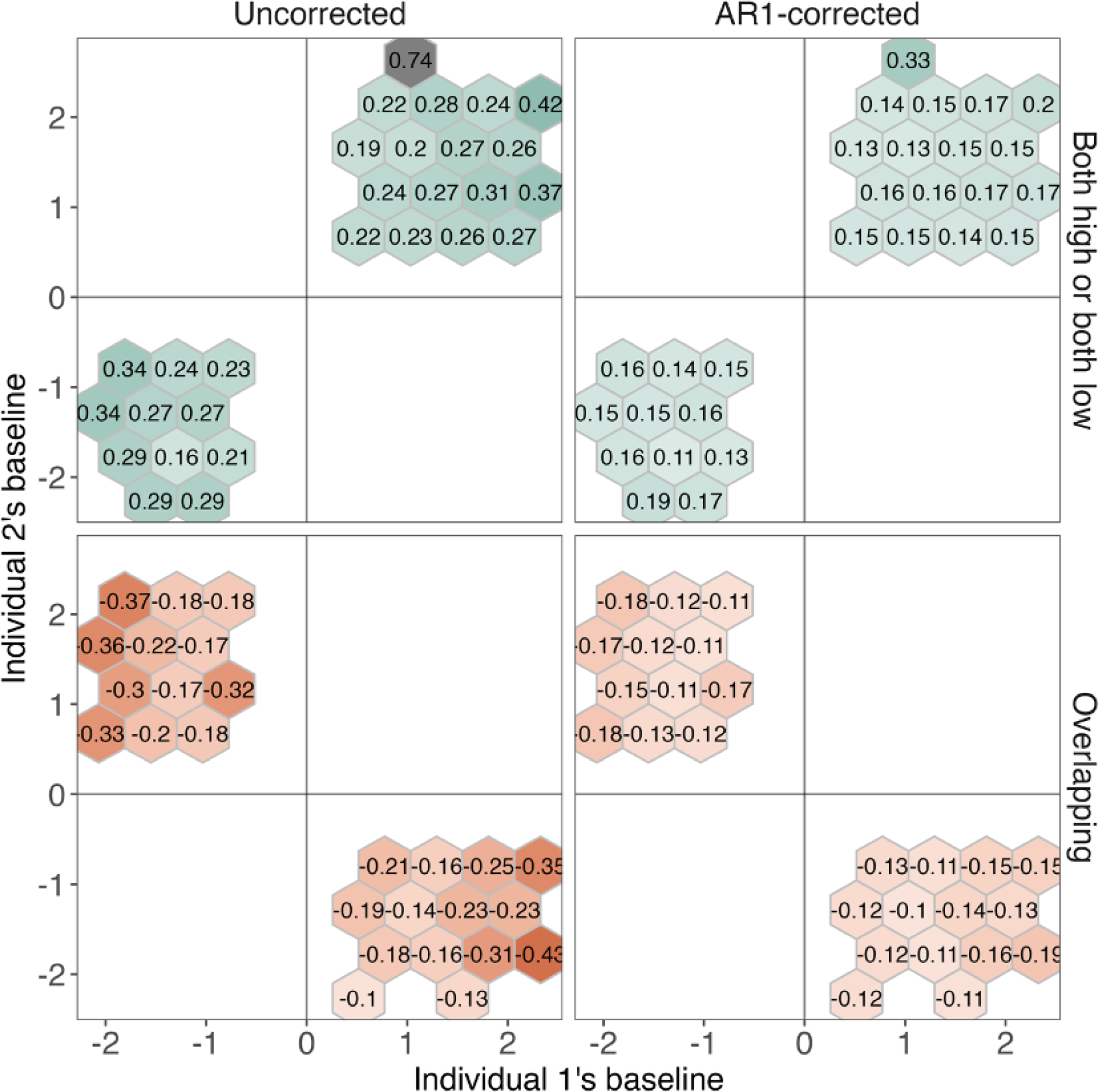
Regression to the mean due to sampling bias in mice. Panel a displays slope estimates for uncorrected (left panels) and AR1-corrected (right panels) models in samples where all pairs are either on the same side (Both high or both low) or different sides of the mean. The x and y axes display the baselines of each individual, scaled within-individual so that baselines can be interpreted as an individual’s distance in standard deviations from their own means. The hexes corresponds to the mean estimated slope for pairs falling within a particular joint baseline area. Colours indicate the magnitude of negative (red) or positive (green) slopes. The central vertical and horizontal lines in each plot represent the “zero” baseline levels for each individual.

**SI Fig. 5.**
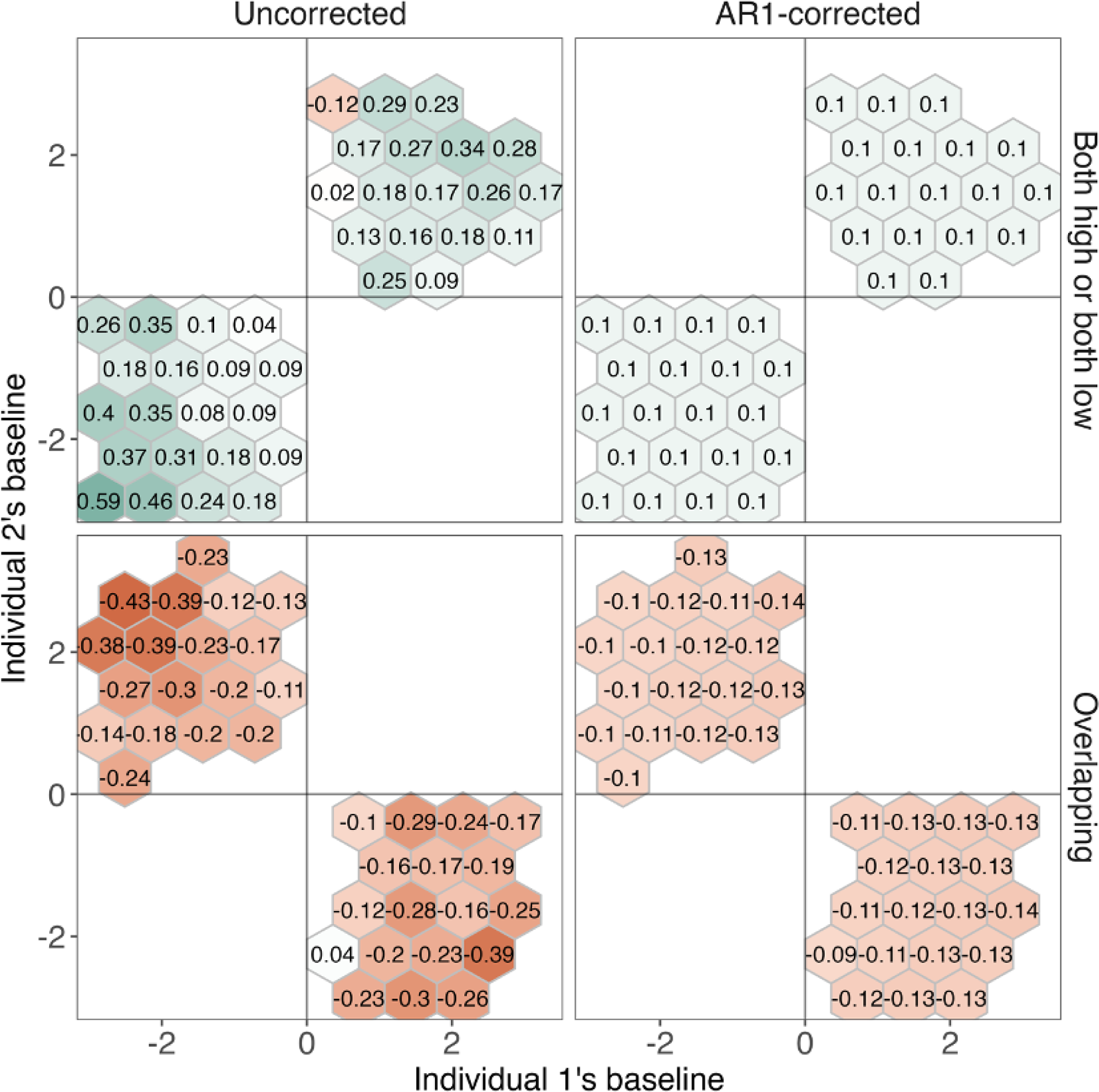
Regression to the mean due to sampling bias in guppies. Panel a displays slope estimates for uncorrected (left panels) and AR1-corrected (right panels) models in samples where all pairs are either on the same side (Both high or both low) or different sides of the mean. The x and y axes display the baselines of each individual, scaled within-individual so that baselines can be interpreted as an individual’s distance in standard deviations from their own means. The hexes corresponds to the mean estimated slope for pairs falling within a particular joint baseline area. Colours indicate the magnitude of negative (red) or positive (green) slopes. The central vertical and horizontal lines in each plot represent the “zero” baseline levels for each individual.

